# Cortical Localization of the Sensory-Motor Transformation in a Whisker Detection Task in Mice

**DOI:** 10.1101/2020.07.08.194555

**Authors:** Behzad Zareian, Zhaoran Zhang, Edward Zagha

**Affiliations:** Department of Psychology, University of California Riverside, 900 University Avenue, Riverside CA 92521 USA; Neuroscience Graduate Program, University of California Riverside, 900 University Avenue, Riverside CA 92521 USA

**Author notes:** equal contributors (Behzad Zareian, Zhaoran Zhang).

## Abstract

Responding to a stimulus requires transforming an internal sensory representation into an internal motor representation. Where and how this sensory-motor transformation occurs is a matter of vigorous debate. Here, we trained mice in a whisker detection go/no-go task in which they learned to respond (lick) following a transient whisker deflection. Using single unit recordings, we quantified sensory-, motor- and choice-related activities in whisker primary somatosensory cortex (S1), whisker primary motor cortex (wMC) and anterior lateral motor cortex (ALM). Based on the criteria of having both strong sensory and motor representations and early choice probability, we identify whisker motor cortex as the cortical region most directly related to the sensory-motor transformation. Our data support a model of sensory amplification occurring between S1 and wMC, sensory-motor transformation occurring within wMC, and propagation of a motor command occurring between wMC and ALM.

## Introduction

To accomplish goal-directed behavior, the brain selects task-relevant stimuli and outputs the appropriate motor responses. A crucial component of this process is the transformation of an internal representation of a sensory stimulus into an internal representation of a motor response. Identifying where this occurs is an essential first step in developing mechanistic understandings of this process. Correlates of sensory-motor transformations in neocortex have been identified in non-human primates (Kim and Shadlen, 1999; Shadlen and Newsome, 2001; de Lafuente and Romo, 2006; Siegel et al., 2015). More recent efforts are now underway to study sensory-motor transformations in mouse neocortex (Matyas et al., 2010; Guo et al., 2014; Yang et al., 2015; Zagha et al., 2015; Goard et al., 2016; Pho et al., 2018; Mayrhofer et al., 2019; Aruljothi et al., 2020; Salkoff et al., 2020), which benefits from less neocortical arealization and the application of novel genetic and physiological tools. Yet, despite these efforts, there is still no agreement as to the location of the sensory-motor transformation.

In this study, we use two major criteria for localizing the site of transformation in mouse cortex. Our first criterion is the coexistence of robust sensory and motor representations. This has been elegantly demonstrated in the primate lateral intraparietal (LIP) cortex during a visual discrimination task; early in the decision process, LIP neurons encode sensory stimulus strength whereas late in the decision process, this activity converges to the anticipated response (Roitman and Shadlen, 2002). Regions with only sensory or only motor representation could be upstream or downstream, respectively, of the transformation process, but cannot mediate the transformation.

Our second criterion is early and robust choice probability (Britten et al., 1996; de Lafuente and Romo, 2006; Crapse and Basso, 2015). Choice probability is a measure of the relationship between neural activity and a behavioral response, independent of stimulus content (Britten et al., 1996). For identical stimulus and behavioral conditions, choice probability is significant only after the initiation of the transformation process. Notable primate studies using multi-site recordings during sensory-motor task performance compared the onset and magnitude of choice probability across multiple cortical regions (de Lafuente and Romo 2006; Siegel, Buschman, and Miller 2015). Regions showing early and robust choice probability are more likely to be initiating the transformation; conversely, regions showing late choice probability are likely reflecting transformations that occurred elsewhere.

We studied a sensory-motor transformation in the context of sensory detection, in which mice learned to respond (lick) following the presence of a transient whisker deflection stimulus. In a recent study using widefield calcium imaging of dorsal cortex, we identified the following regions as potentially contributing to the transformation process by expressing robust activity between stimulus onset and response: whisker representation of primary somatosensory cortex (S1), whisker region of primary motor cortex (wMC), and anterior lateral motor cortex (ALM) (Aruljothi et al., 2020). Previous studies of similar sensory-motor pairings (whisker stimulus→lick) provide partial support for the transformation occurring within each region. S1 shows robust sensory encoding (Stüttgen and Schwarz, 2008; O’Connor et al., 2010; Wang et al., 2012; Sachidhanandam et al., 2013), can evoke motor responses (Matyas et al., 2010), and displays significant choice probability (Sachidhanandam et al., 2013; Yang et al., 2015; Chen et al., 2016; Kwon et al., 2016). wMC shows robust sensory and motor encoding (Ferezou et al., 2007; Huber et al., 2012; Zagha et al., 2015) and displays neural dynamics consistent with linking a sensory stimulus to a motor response (Zagha et al., 2015). ALM shows robust motor encoding (Li et al., 2015; Chen et al., 2017) and displays neural dynamics consistent with motor planning (Inagaki et al., 2018). Moreover, acute perturbation of all three regions impairs whisker detection (Huber et al., 2012; Guo et al., 2014; Yang et al., 2015; Zagha et al., 2015). However, previous studies have not compared sensory-, motor-, and choice-related content across all three regions in the same task. Moreover, it is critical that such studies are conducted with sufficient temporal resolution to determine the precise timing of these signals in each region.

In this study we measured single unit activity in S1, wMC and ALM during a whisker detection task. Based on analyses of sensory and motor encoding and choice probability, we find that activity in wMC is most correlated with a sensory-motor transformation process.

## Materials and Methods

### Subjects

Animals and experiments were approved by the IACUC of University of California, Riverside. Both male and female, adult mice were used in the experiments, from genetic backgrounds C57BL/6J and BALB//cbyJ. The mice were kept in 12 hours light and dark cycle and the experiments were conducted during the light cycle.

### Surgery

Mice were anesthetized using an induction of ketamine (100 mg/kg) and xylazine (10 mg/kg) and maintained under isoflurane (1-2%) anesthesia. A 10×10 mm portion of the scalp was removed and a lightweight metal headpost was attached to the skull using cyanoacrylate glue. The headpost includes an 8×8 mm central window, leaving the skull over dorsal cortex exposed. The exposed skull was sealed with a thin layer of cyanoacrylate glue and covered with silicone gel. They were further treated with meloxicam (0.3 mg/kg) and enrofloxacin (5 mg/kg), the day of the surgery and for 2 additional days after the surgery. After recovery from surgery for a minimum of three days, water restriction was initiated and the mice were introduced to the behavioral task.

### Behavior

Matlab software and Arduino boards were used to control the behavioral task flow. The mice were head-fixed in the setup during a behavioral session. Piezoelectric benders with attached paddles were placed within the whisker fields bilaterally. One side was assigned as target and the other as distractor at the onset of training and remained consistent throughout training and recording. The location of the paddles was in the mid-ventral whisker fields (targeting D2/E2-D3/E3 whiskers), with movement in the caudal direction of 1 mm for our largest stimuli. A voltage generator (ThorLABS) was used to drive the piezo benders. Whisker deflections were triangular waves of 2 to 200 ms. Amplitude and velocity of deflections were varied to operate within the dynamic range of each mouse’s psychometric curve. In any single recording session, two stimulus amplitudes were applied: one near the saturation of the psychometric curve and one 2x or 4x lower within the dynamic psychometric range. For each session, equal strength stimuli were presented for target and distractor trials. Licking responses were detected by an infrared beam break circuit positioned immediately in front of a central lickport. Reward was approximately 5 μL of water. Mice were trained in three stages, progressing from 1) classical condition to 2) operant conditioning to 3) the full task with punishment for incorrect responses (see (Aruljothi et al., 2020) for training details and learning trajectories). Intertrial intervals (ITI) varied from 6 to 10.5 seconds, drawn from a decreasing exponential distribution, to correct for an expectation hazard function and thereby minimize a timing strategy (Elithorn and Lawrence, 1955). Trial types consisted of target trials (deflection of the target paddle), distractor trials (deflection of the distractor paddle) or catch trials (no stimulus). The initial percentages of each trial-type were set as follows: target trials 15%, distractor trials 60%, catch trials 25%. Immediately following stimulus onset was a lockout period of 200 (46 sesssions) or 300 ms (8 sessions). Licking during the lockout period resulted in aborting the current trial. Following the lockout period was a 1 second response window. Responses within the response window following target stimuli (hits) were rewarded with a fluid reward. Withholding (not responding) on a distractor trial was rewarded with a shortened ITI (1.4 to 3.1 second distribution) and subsequent target trial. In expert mice, the percentage of target and distractor trials were similar across each session. All licking outside the post-target response window (including during the ITI) was punished by resetting the ITI. Behavioral sessions typically lasted between one to two hours, which included 200 to 400 trials. Mouse weights were maintained above 85% of their initial weights by either receiving all the water from task or receiving additional water and wet food after the task.

### Engagement period

For behavioral and recording analyses, the trials were truncated to engaged periods using a gap of 60 seconds as a disengagement criterion. For each session, the single, longest continuous bout was used for further analyses. Sessions without continuous engagement for 10 minutes were excluded from further analysis. We also excluded the trials in which the mice responded prematurely (licking during the lockout period).

### Behavioral analysis

For behavioral metrics, hit rate was obtained by dividing the number of hits by the total number of target deflection trials. Spontaneous rate was obtained by dividing the number of spontaneous responses by the total number of catch trials. For the sessions that did not include catch trials, the 1 second pre-stimulus response rate was used as a replacement of the spontaneous rate. For the purpose of d-prime calculations, response rates of 0 and 1 were estimated at 0.01 and 0.99. Behavioral d-prime was calculated as:

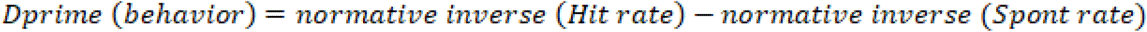

Mice were considered expert in the task once they achieved a d-prime > 1 for three consecutive days. Mice were recorded immediately following acquisition of expert status. For behavioral performance measures during the electrophysiological recording sessions, see Figure 1D.

**Figure 1.**
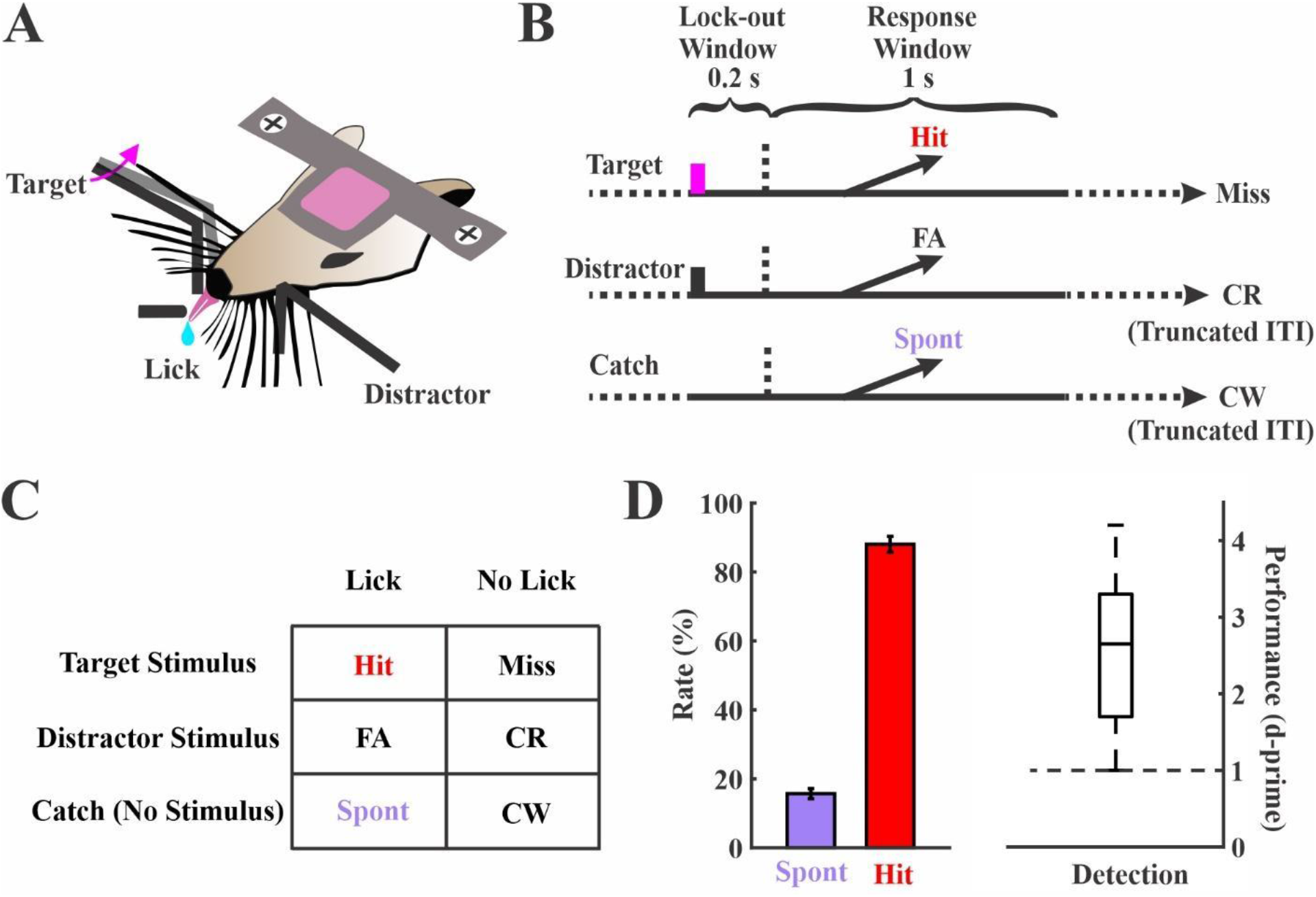
Sensory detection task structure and performance. (**A**) A side-view of the task showing bilateral paddle placement and the central lickport. Head-fixed mice learned to respond to whisker deflection on one side (target) by licking the central lickport to obtain a fluid reward, and to ignore the deflections on the contralateral side (distractor) by withholding a licking response. (**B**) Trial structures. Each trial starts either with a target deflection (magenta bar, target), a distractor deflection (black bar, distractor) or no stimulus (catch). Responding during the lockout window (indicated by the horizontal dashed lines) aborted the current trial. (**C**) Possible outcomes based on trial type and response: Hit, Miss, False Alarm (FA), Correct Rejection (CR), Spontaneous Response (Spont) and Correct Withholding (CW) (**D**) Performance of all the 54 sessions that were included in this study collected from 19 expert mice. Boxplot for d-prime values shows min, max, median and 25^th^ and 75^th^ percentiles.

### Electrophysiology

Craniotomies and durotomies of less than 0.5 mm in diameter were established on the day of recording, under isoflurane anesthesia. After 30 to 60 minutes post-surgery, mice were tested in the behavioral task without electrode implantation to ensure recovery to normal behavior. Upon evidence of normal expert behavior, a silicon probe (Neuronexus A1×16-Poly2-5mm-50s-177) was advanced into the brain using a Narishige micro-manipulator under stereoscope guidance. We positioned the recording sites to target layer 5, from 500 to 1000 um below the pial surface. Recording sites were targeted to the barrel field of primary somatosensory cortex (S1), the whisker region of primary motor cortex (wMC) and anterior lateral motor cortex (ALM), based on the functional mapping studies of (Aruljothi et al., 2020). Precise coordinates (mm, from bregma): S1 3.2-3.7 lateral, 1-1.5 posterior; wMC 0.5-1.5 lateral, 1-2 anterior; ALM 1-2 lateral, 2-2.5 anterior.

Whisker alignment for S1 recordings was verified by three methods. First, after electrode implantation we verified correct alignment by hand mapping of several individual whiskers and observing LFP responses. Second, we only included sessions with clear peaks in the combined multi-unit post-stimulus time histogram. Third, we only included sessions with larger target encoding than distractor encoding across the population (1.5x greater variance for target versus distractor d-prime). In contrast, inclusion of wMC and ALM sessions were based solely on anatomical location.

### Electrophysiology pre-processing and spike sorting

Neuralynx software was used for data acquisition and spike sorting. Electrophysiological signals were sampled at 32 kHz; wideband signals were band-pass filtered from 0.1Hz to 8000 Hz and signals for spike sorting were additionally high pass filtered at 600 Hz to 6000 Hz. Putative spikes were identified as threshold crossings over 20 to 40 μV, set at the beginning of each recording session to be well isolated from baseline noise. Spike sorting and clustering was done offline using KlustaKwik algorithm in SpikeSort3D software. The clusters were further manually inspected and merged based on the similarity of waveform and cluster location in peaks and valleys feature space; clusters indicative of movement artifacts (non-spike waveform, equal amplitude in all channels) were removed. Additionally, we used isolation distance (ID) and L ratio to verify cluster quality (mean +/− standard error of mean (SEM): ID 15.6180+/− 0.7378, L ratio 0.2312+/− 0.0172). Further data analyses were conducted using MATLAB software (Mathworks). Spike times were binned within 5 ms non-overlapping bins. Reaction time binning in Figure 2 used the following bins: (ms) Sensory: fast RT 201-249, medium RT 250-330, slow RT 335-547; Sensory-Motor: fast RT 322-374, medium RT 374-447, slow RT 459-982; Motor: fast RT 301-326, medium RT 329-399, slow RT 407-1240.

**Figure 2.**
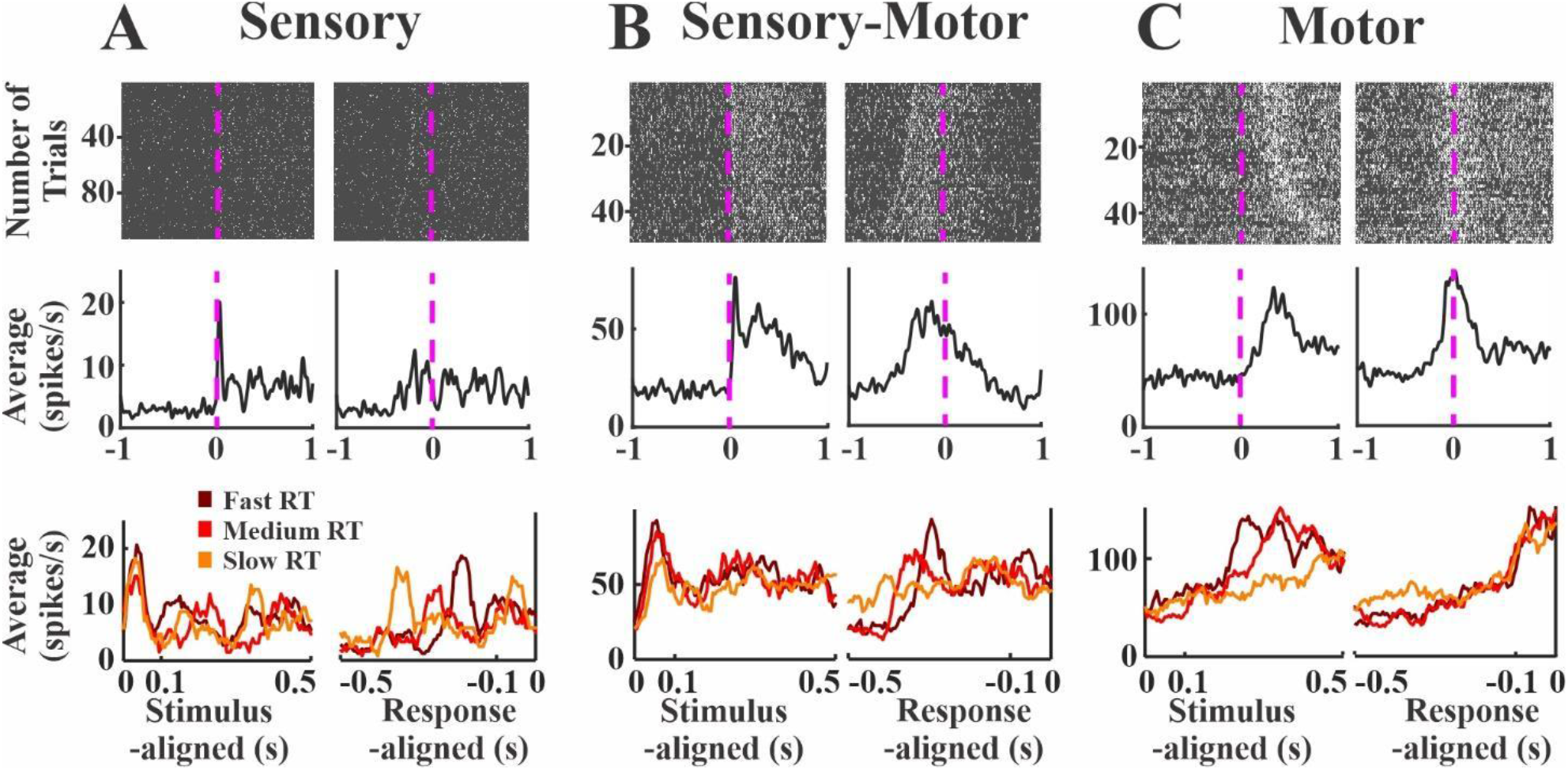
Examples of sensory, sensory-motor and motor units. (**A**) A sample sensory unit from S1. (Top) Raster plots show spiking activity for all trials within a session, aligned to the stimulus onset (left) and the mouse’s reaction time (right). Stimulus and response events are indicated by the magenta dashed lines. The trials in all raster plots are sorted according to the mouse’s reaction time. (Middle) Average spiking rates across all trials. A transient peak immediately post-stimulus is observable with sensory alignment (left) but not with motor alignment (right). (Bottom) Trials were further grouped into slow, medium and fast reaction times. The sensory peak overlaps in all groups when aligned to the stimulus onset (left) but varies when aligned to the reaction time (right). (**B**) Same structure as panel [A], but for a sample sensory-motor unit in wMC. (Middle) A transient sensory peak is observable with sensory alignment (left), along with a ramping activity prominent in the motor alignment (right). (**C**) Same structure as panel [A] but for a sample motor unit in ALM. (Middle) Motor alignment shows a prominent ramping immediately prior to the reaction time. (Bottom) Unlike the sensory unit, the stimulus-aligned peak activity varies with reaction time.

### Sensory encoding

Sensory encoding was quantified using a neurometric approach based on signal detection theory that enables the direct comparison of neural performance to behavioral performance (Britten et al., 1992; Stüttgen and Schwarz, 2008). For this analysis, the target and distractor trials were used regardless of their outcome (hits, misses, false alarms and correct rejections). Data presented for sensory encoding used the larger of the two stimuli for neurometric and psychometric comparisons. ‘Stimulus present’ data were spike counts within 100 ms immediately post-stimulus; ‘stimulus absent’ data were spike counts within three consecutive 100 ms epochs pre-stimulus onset. Distributions based on single trials were compared using receiver operating characteristic (ROC) curve, by plotting the cumulative distribution function of each distribution against each other. The area under the ROC (AU-ROC) was converted to neurometric d-prime as:

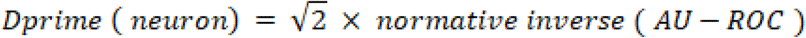

AU-ROC was bounded by 1-0.003 for population encoding, to ensure the output of real numbers.

### Combining units

In figure 4, sensory encoding was calculated not only for single units, but also for different combinations of units. In figure 4B set 3, the spikes were summed together for each 5 ms bin across all the units recorded in a session. This results in a single multi-unit per session, for which sensory encoding was calculated similar to single units. In figure 4B set 4, the spikes of all units in each region were combined across all sessions. To equalize the number of trials across sessions, the sessions with less trials had their trials multiplicated and added after the original trials to match the trial number of the session with the most trials; for the sessions with the trial numbers not a common divisor of the trial number in the longest session, the trials were randomly sampled (with replacement) from these sessions accordingly and added to that session to fill in. The sensory encoding for these combined units and trials were calculated similar to the previous cases.

### Random sampling

In addition to combining all units from each session or region, we ran additional analyses to assess encoding for random sets of units (Figure 4C and D).We randomly selected units to be added sequentially and computed d-prime values for each group, with group size spanning 1 to total number of units per region. We permutated this ordering and d-prime calculation 300 times and plot the mean +/− standard deviation curve in Figure 4C. For the purpose of neurometric-psychometric comparison, we transposed the data by creating a histogram in which each bin is a specific d-prime value (bin width of 0.02 spanning 0 to 4.5) and the entries (dependent variable) are the number of units with a combined d-prime value within each bin. Mean +/− standard deviation for the number of combined units to achieve a specific d-prime is plotted in Figure 4D.

### Sensory-motor alignment

A common method used to assess sensory and motor content is to determine the temporal alignment of neural activity to stimulus and response onsets (Hanes and Schall 1996; Mountcastle et al. 1974; Romo et al. 2002). To quantify sensory and motor content, we used a similar neurometric approach as described above. Because the motor alignment requires responding to the stimuli, we only considered hit trials in this analysis. For sensory alignment, we used the same 100 ms post-stimulus window as for sensory encoding. For motor alignment, we used a 100 ms pre-response window. Both conditions were compared to the same pre-stimulus baseline as described above.

### Latency estimation

Latencies of activation after the stimulus onset was estimated by using a 20 ms sliding window (75% overlap) post-stimulus, comparing to a pre-stimulus baseline, for all target trials. Baseline activity was the average activity in 20 ms during the 1 second pre-stimulus epoch. We excluded the first 10 ms after the stimulus onset due to possible contamination with stimulus artifacts.

### Choice probability

Choice probability was calculated as the separation of neural activity on hit versus miss trials. All spikes from each session were combined to increase spike density for comparisons. To ensure an adequate number of trial types and ensure valid comparisons: 1) we calculated the hit rate for small and large amplitude stimuli separately 2) if the difference between those was below 15%, trials from both types of stimuli were pooled together 3) if the difference was above 15%, the stimulus type with larger number of hits and misses were considered 4) all sessions with fewer than 5 misses were removed. We used the AU-ROC method along sliding time windows to calculate choice probability as the separation between spiking distributions on hit vs miss trials. The duration of the sliding window was set to 50 ms with 90% overlap.

### General Statistics

We used permutation statistics for comparing sensory-motor variance and slope differences (10,000 repetitions). Briefly, we shuffled the units between the conditions (for instance, S1 and wMC sensory encoding d-primes), and we pooled two new putative sets and calculated the difference in variable of interest (for instance, variance). Then we assessed the position of the actual variance difference among these 10,000 repetitions and we reported the p-value as the proportion of the repetitions above the actual variance (two-sided calculation). For comparing random sampling results, Cohen’s d was used by dividing mean difference of the two groups by their pooled standard deviation. For calculating significant choice probability within each region across sessions, for each time window, we calculated a one sample t-test between the reported choice probability and chance level (0.5) (p-value=0.01). For comparing choice probability amplitudes across regions, we used unpaired t-tests with an alpha level corrected for multiple comparisons with the Bonferroni method (0.05/3). For latency estimation, paired t-test was used (between each 20 ms window and baseline) with alpha level of 0.05. All of the bars (shaded or whisker) show mean +/− SEM unless otherwise indicated.

## Results

### Behavioral task and electrophysiological recordings

Head-fixed mice were trained to perform a whisker detection go/no-go task in which they learned to lick a lickport following a transient whisker deflection in one whisker field (target) to obtain a fluid reward (Figure 1A). Stimuli were piezo-controlled caudal deflections of a paddle contacting multiple whiskers. We imposed a minimum lockout period of 200 between stimulus onset and response window to separate sensory from motor encoding. Trials were aborted if any responses occurred during the lockout period. Target trials were interleaved with two other trial types: distractor trials, in which there was a transient deflection of the same amplitude in the opposite whisker field, and catch trials, in which there was no stimulus deflection (Figure 1B and 1C). Mice were considered expert in this task once they achieved a detection d-prime (separation between hit rate and spontaneous rate) greater than 1 for three consecutive days. Electrophysiological recordings were conducted in expert mice while performing the detection task. For the recording sessions included in this study, the behavioral performance measures: hit rate 88.0% +/− 2.3%; spontaneous rate 15.7% +/− 1.4%; d-prime 2.6 +/− 0.1 (n=54 sessions from n=19 mice, Figure 1D).

Based on a concurrent widefield calcium imaging study (Aruljothi et al., 2020), we targeted our electrophysiological recordings to three cortical regions contralateral to the target whisker field: whisker representation of primary somatosensory cortex (S1), whisker region of primary motor cortex (wMC) and anterior lateral motor cortex (ALM). Each of these regions were significantly active post-stimulus and pre-response (Aruljothi et al., 2020), and therefore may contribute to the sensory-motor transformation process. We used silicon probes with contact sites spanning layer 5 to record multiple single units in each region (S1: 445 units, 25 sessions; wMC: 424 units, 16 sessions; ALM: 315 units, 13 sessions). To establish the functional hierarchy of these regions, we calculated post-stimulus response latency for each session. Latency measurements are consistent with the functional ordering of S1→wMC→ALM (mean +/− standard deviation: S1 30 +/− 15 ms, wMC 48 +/− 28 ms, ALM 95 +/− 43 ms; ANOVA F(2,51)=23.56, p<0.01; Tukey post-hoc comparison: S1-ALM p<0.01, wMC-ALM p<0.01, S1-wMC p=0.11).

### Sensory vs motor representation

The next set of analyses quantify the sensory and motor content within each cortical region. To illustrate how we distinguish ‘sensory content’ from ‘motor content’, we present three example neurons in Figure 2 with robust sensory, sensory-motor, and motor context, respectively. The ‘sensory’ unit shows robust alignment to the stimulus onset as a sharp peak in the average spiking activity across trials (Figure 2A left column). In contrast, this unit lacks a sharp peak in average activity when aligned to the response (Figure 2A, right column). This is further apparent when grouping the trials based on reaction times (Figure 2A, bottom row): peak activity levels overlap regardless of reaction time when aligned to stimulus onset, whereas peak activity levels vary according to reaction time when aligned to the response. On the other hand, the ‘motor’ unit shows prominent alignment to the response (Figure 2C, right column) with activity that is delayed when aligned to the stimulus onset (Figure 2C, right column). In further contrast with the ‘sensory’ unit, peak activity levels in the ‘motor’ unit overlap when aligned to the response but not to the stimulus onset (Figure 2C, bottom row). The ‘sensory-motor’ unit shows a mixture of both features, with sharp, transient activity aligned to the stimulus followed by activity that is sustained until the response (Figure 2B). Note that the time-locked ‘sensory’ responses occurred within the first 100 ms post-stimulus and the time-locked ‘motor’ responses occur within the last 100 ms pre-response. These time windows were used for the analyses below.

### Quantification of sensory encoding

Figure 3A shows the average spiking activity for an example unit from target-aligned S1, contralateral to the target whisker field. On target trials (Figure 3A blue), this unit displayed a prominent increase in spiking immediately after stimulus presentation followed by a lower level of persistent activity. Spiking activity on distractor trials (Figure 3A, black), in contrast, appeared only slightly elevated from pre-stimulus levels. In order to quantify stimulus encoding, we used the neurometric d-prime approach (Figure 3B-E), which accounts for single trial variability and allows for comparisons between neuronal performance and behavioral performance (Britten et al., 1992). We compared trial by trial distributions of pre-stimulus and post-stimulus spiking activities (Figure 3B and 3C). For the post-stimulus condition, we included spikes in the first 100 ms post-stimulus onset. We calculated the d-prime value of each unit from area under the curve (AUC) of the receiver operating characteristic (ROC) function (AU-ROC) between the pre-stimulus and post-stimulus distributions (Figure 3D). For this analysis, a d-prime greater than zero indicates higher spiking activity post-stimulus compared to pre-stimulus. Plotting the d-prime values for all units in this S1 recording session (Figure 3E) shows target versus distractor stimulus encoding across the population. As shown in this example session, target stimulus encoding is highly variable yet positively skewed across these units, whereas distractor stimulus encoding is considerably more restricted.

**Figure 3.**
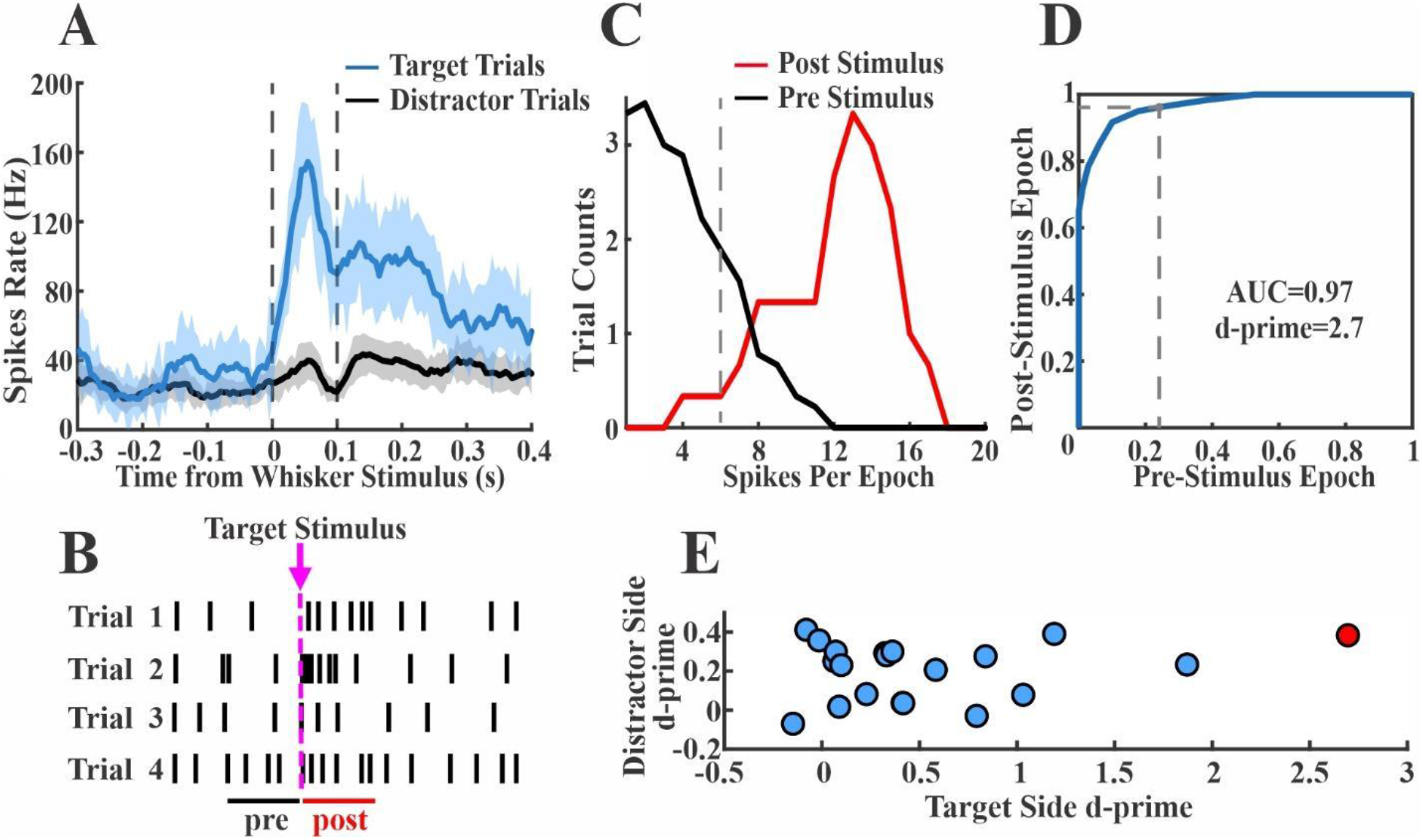
Quantification of target and distractor stimulus encoding. (**A**) A sample S1 unit firing rate averaged across target (blue) and distractor (black) trials. This unit shows a prominent increase in spiking after target stimulus onset. Dashed lines reflect the post-stimulus window used for quantification of sensory encoding. (**B**) Illustration of the single trial pre-stimulus and post-stimulus windows. (**C**) Plot of pre-stimulus and post-stimulus spike count distributions from target trials of the example unit shown in [A]. (**D**) Plotting of the pre-stimulus and post-stimulus cumulative distribution functions to create a receiver operating characteristic curve for the example unit shown in [A]. The area under the curve is transformed into a neurometric d-prime value. The large response in [A] is reflected in the large separation of pre-stimulus and post-stimulus distributions in [C] and the highly convex ROC curve in [D]. (**E**) Scatter plot of all single units in this recording session, plotting target d-prime vs distractor d-prime values (example unit indicated in red). Note that target d-prime values are more positively skewed than distractor d-prime values.

Figure 4A shows target and distractor stimulus encoding for S1, wMC and ALM across all recorded neurons, indexed to the average behavioral performance of the mice during the corresponding recording sessions. Both S1 and wMC neurons showed prominent target stimulus encoding across their populations. ALM neurons, in contrast, showed minimal target stimulus encoding. We analyzed these data with both single unit and population approaches (Figure 4B-D). First, we compared the group means of single unit target encoding across these three regions (Figure 4B set 2). We found mean target stimulus encoding to be significantly higher for S1 and wMC compared to ALM, and, interestingly, for wMC to be significantly higher than S1 (S1: 0.23 +/− 0.02; wMC: 0.37 +/− 0.03; ALM: 0.01 +/− 0.01; ANOVA F(2,1181) = 51.80, p <0.01; Tukey post-hoc comparison: S1-ALM p<0.01, wMC-ALM p<0.01, wMC-S1 p<0.01; effect size: wMC 61% larger than S1). Additionally, we compared the summed spiking from multiple neurons in each trial (summed within each recordings session, Figure 4B set 3, and summed across all units within each region, Figure 3B, set 4). When combined across each population, the neurometric d-prime for S1 and wMC, but not ALM, outperformed the behavioral d-prime.

To quantify population coding within each region, first we randomly sampled different numbers of units in each region and plotted the resulting neurometric target d-prime values (see Methods) (Figure 4C). As the number of sampled units increased, the target d-prime increased well beyond behavioral performance for S1 and wMC, but not for ALM. Furthermore, this trend rose faster for wMC than for S1 (Figure 4C). Next, we transformed the data across axes (Figure 4D). This allowed us to quantify how many units in each region were required to match behavioral performance. Both S1 and wMC populations were able to match behavioral performance (Figure 4D, red arrows), whereas the ALM population was not. Moreover, fewer units were required to match behavioral performance for wMC compared to S1 (mean +/− standard deviation: S1 95 +/− 44 units; wMC 52 +/− 25 units; Cohen’s d = 1.2). Altogether, these results demonstrate robust sensory encoding in S1 and wMC but not in ALM, with increased sensory encoding in wMC compared to S1.

**Figure 4.**
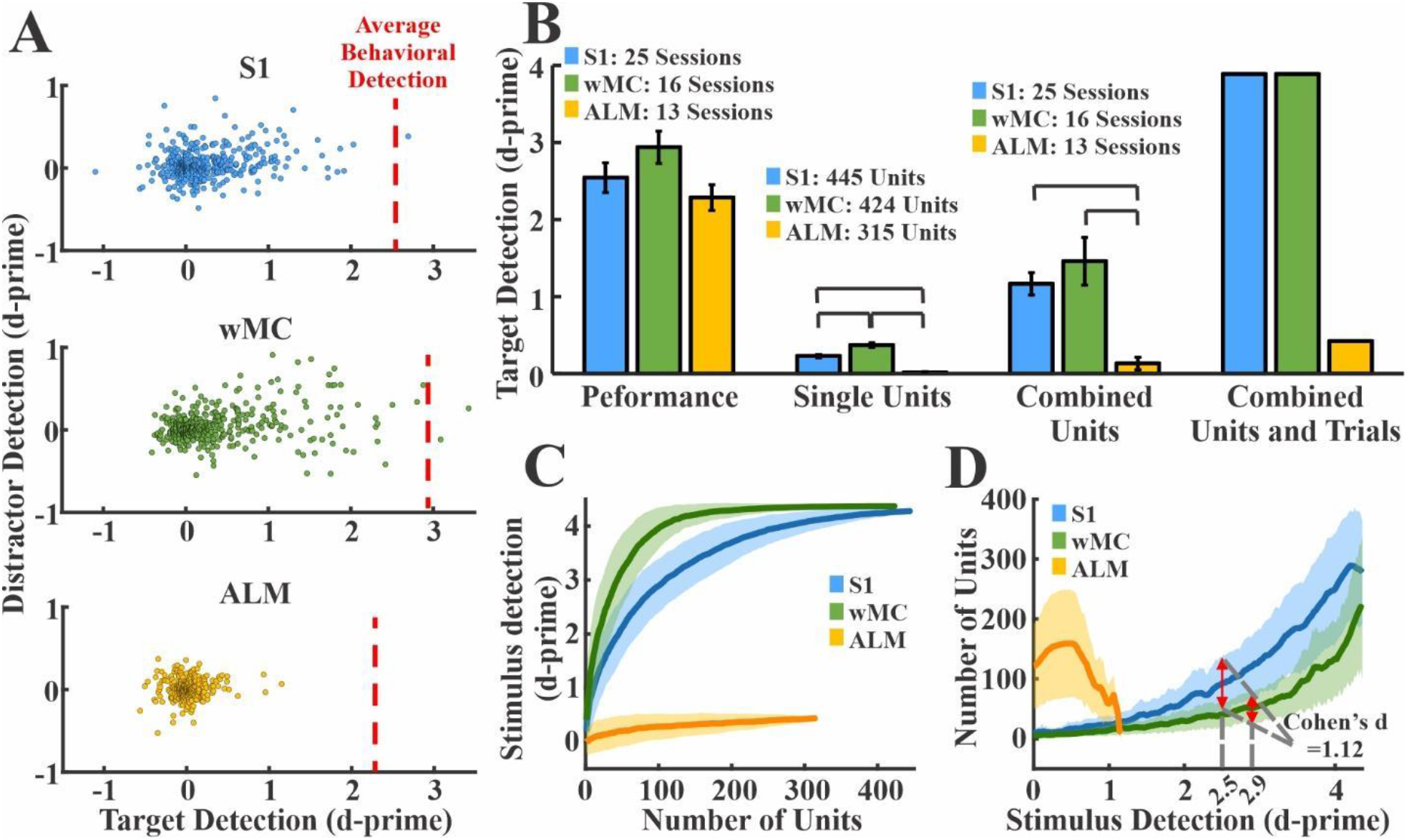
Sensory encoding and neurometric-psychometric comparisons across cortical regions. (**A**) Distribution of single unit target and distractor d-prime values for all S1 (top, blue, n=445 units), wMC (middle, green, n=424 units), and ALM (bottom, yellow, n=315 units) units. The average behavioral detection performance (behavioral d-prime) during these recording sessions is depicted by the red dashed lines (S1 = 2.5, wMC = 2.9 and ALM = 2.2). Note that S1 and wMC target d-prime values are highly positively skewed along the x-axis (target detection) but ALM units are not. (**B**) Behavioral and neural d-prime measures across regions. Lines connecting columns within each set denote differences of statistical significance. Set 1, psychometric d-prime across all regions. Set 2, neurometric d-prime averaged across all single units within each region. Set 3, neurometric d-prime of summed spiking within each session averaged across all sessions. Set 4, neurometric d-prime of summed spiking of all units within each region. Combining units results in neurometric performance surpassing psychometric performance for S1 and wMC, but not ALM. (**C**) Random combinations of units and assessment of resulting neurometric d-prime values. Increasing the number of combined units increased d-prime values, with the fastest rate of rise in wMC. (**D**) Transformation of data in panel [C], reflecting the number of combined units achieving the corresponding d-prime values. Red arrows overlaying S1 and wMC data indicate the number of units needed to match behavioral performance. Fewer wMC units were required to match behavioral performance compared to S1 and ALM. The traces and shades in panels [C] and [D] are the mean +/− SD across 300 iterations.

### Sensory and motor alignments across cortical regions

Next, we sought to assess the sensory versus motor alignment across these three cortical regions. For quantification, we used a similar neurometric d-prime method as above, yet for only hit trials and for both sensory and motor alignments (Figure 5). For sensory alignment, we again analyzed spiking within 100 ms following stimulus onset; for motor alignment, we analyzed spiking within 100 ms preceding the reaction time (Figure 5A). Due to our imposed lockout between stimulus onset and response window, these analysis epochs did not overlap. In Figure 5B, we plot the sensory vs motor alignment for each unit across all three regions. Interestingly, much of the variance of the S1 and wMC populations lies along the diagonal, indicating equal sensory and motor alignment in these regions. In contrast, the variance of the ALM population is largely along the x-axis, indicating predominant motor alignment. We quantified this by calculating a sensory-motor variance ratio: variance along sensory axis divided by variance along motor axis. Indeed, this variance ratio was similar for S1 and wMC, and both were significantly larger than ALM (variance ratio: S1 = 0.90, wMC = 0.86, ALM = 0.09; permutation statistics, S1-wMC, p=0.79, S1-ALM, p<0.01, wMC-ALM, p<0.01). We also used the variance of each alignment as a measure of representation and compared these values within and between regions (Figure 5C). Similar to the findings depicted in figure 3, we found that sensory alignment increased from S1 to wMC, then fell dramatically in ALM. Additionally, we found that motor alignment increased from S1 to wMC and ALM, with similar variance in wMC and ALM.

**Figure 5.**
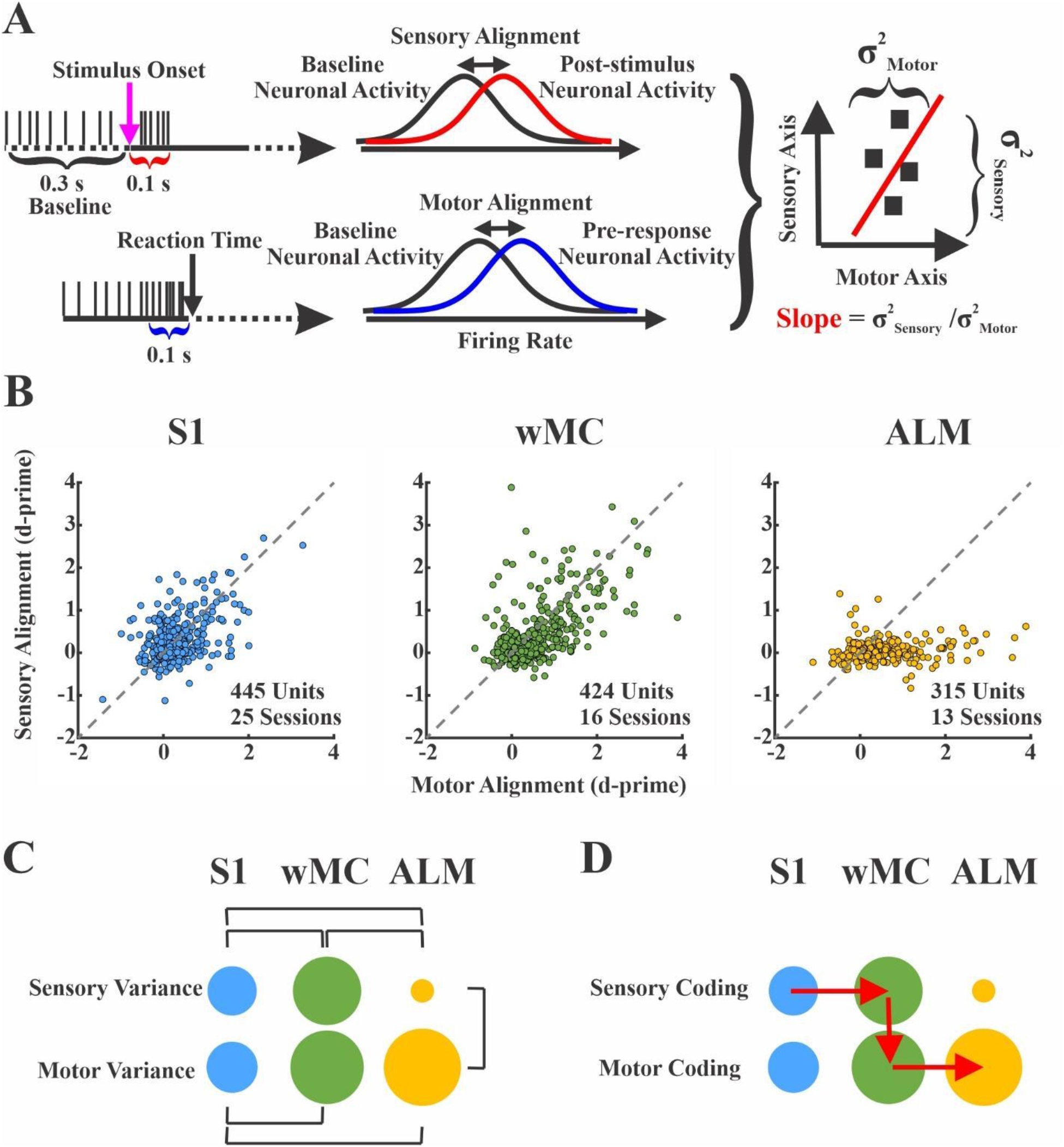
Sensory and motor representations on hit trials across cortical regions. (**A**) A schematic showing how sensory-motor alignment was calculated. 100 ms windows, after stimulus onset (magenta arrow) and preceding time reaction time (black arrow), were referenced as sensory (red) and motor (blue) epochs, respectively. Spike counts in these windows were compared to a pre-stimulus baseline (black). (Right) Sensory-aligned vs motor-aligned values were plotted for each unit. Population measurements of each region included the sensory and motor variance (sigma squared) and slope (sensory variance / motor variance). (**B**) Sensory and motor alignment for all of the recorded units of S1 (left, n=445), wMC (middle, n=424) and ALM (right, n=315). In each plot, the x-axis and the y-axis show motor and sensory alignment d-prime values, respectively. The dashed line indicates equal sensory and motor alignment. Note that S1 and wMC populations both show high variance along the unity line, whereas the ALM population shows high variance nearly exclusively along the motor-aligned axis. (**C**) Each circle’s area is proportional to the variance along the indicated axis. Statistically significant differences are indicated by bars (permutation statistics). Note the increase in both sensory and motor variance from S1 to wMC and reduction in sensory variance in ALM. (**D**) A proposed sensory-motor propagation pathway (red arrows), indicating sensory propagation between S1 and wMC, sensory-motor transformation within wMC, and motor propagation between wMC and ALM.

The above analyses support the observation that S1 and wMC show both sensory-aligned and motor-aligned content, and therefore meet our first criterion for identifying the location of the sensory-motor transformation. ALM, in contrast, shows only motor-aligned content, which we interpret as being downstream of the transformation process. Based on the magnitudes of sensory and motor alignment we propose the model in Figure 5D, with the sensory-motor transformation occurring within wMC.

### Choice probability across cortical regions

To further test this model, we applied an additional analysis that does not rely on temporal alignments. Here, we calculated choice probability across time for each of the three regions. Choice probability quantifies the separation between hit and miss trials, thereby isolating response-related activity (Britten et al., 1996; de Lafuente and Romo, 2006; Crapse and Basso, 2015). According to our second criterion, the region with early and robust choice probability is most likely to initiate the sensory-motor transformation. For these analyses we combined spikes from all units within each recording session to enhance spike density per comparison. Figure 6 shows the average spiking activity on hit and miss trials from three example sessions. All sessions here show higher activity on hit trials during some portion of their post-stimulus response, indicating positive choice probability. However, there are notable differences between sessions. Both the S1 and wMC sessions show increased activity on hit trials immediately post-stimulus and during the response window. However, the difference appears to be larger and more sustained for the wMC session. In contrast, the ALM session shows a robust peak selectively on hit trials, which emerges gradually after stimulus onset.

**Figure 6.**
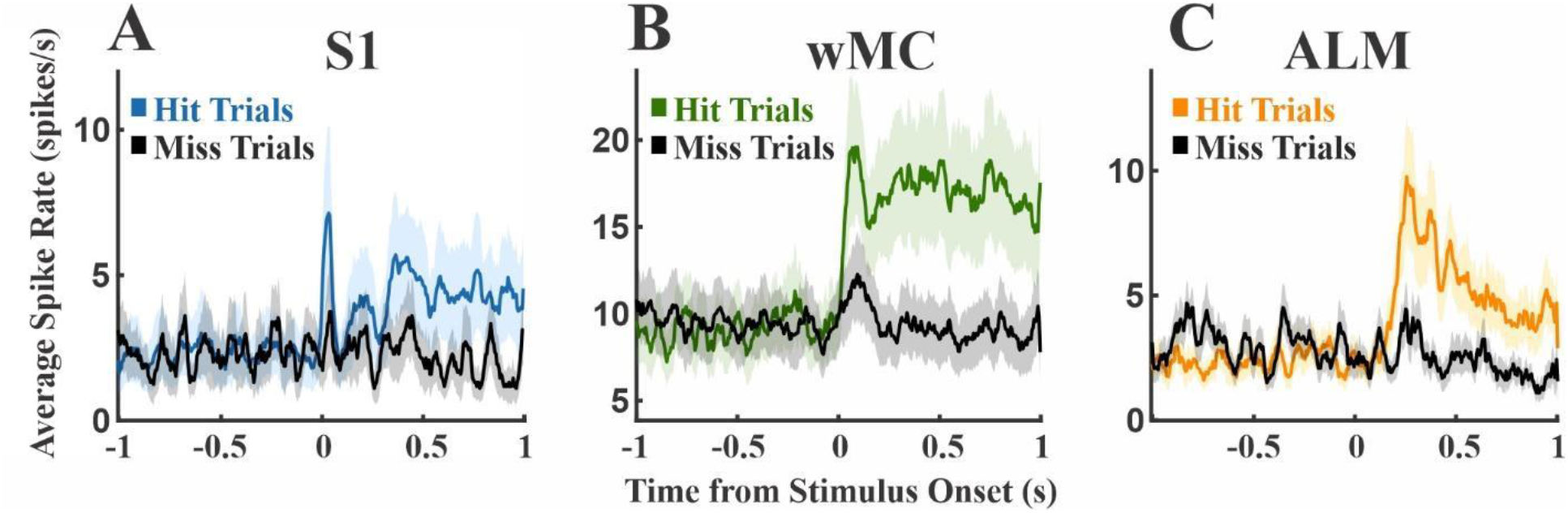
Comparison of spike rates on hit versus miss trials in example recording sessions. Colored plots denote hit trials, black plots denote miss trials. (**A**) An example S1 session, showing moderately higher hit-related spiking immediately post-stimulus and during the response window. (**B**) An example wMC session, showing robust increased and sustained hit-related spiking that emerges immediately post-stimulus. (**C**) An example ALM session, showing robust increased hit-related spiking that emerges late post-stimulus.

We calculated choice probability in 50 ms sliding windows across sessions for each region (Figure 7A). All three regions showed significant increases in choice probability post-stimulus (Figure 7B, gray bars; one-sample t test, comparing to chance level at 50% and alpha level of 0.01), indicating higher spiking rate on hit trials. Interestingly, S1 additionally showed significant negative choice probability pre-stimulus (Figure 7B, purple bars), indicating that lower spike rates immediately before the stimulus onset predicts a hit response. For all three regions, significant post-stimulus choice probability preceded the reaction time, which was always >200 ms due to our lockout period. Yet, significant post-stimulus choice probability emerged earliest in wMC compared to S1 and ALM (S1 165 ms, n=21 sessions; wMC 70 ms, n=13 sessions; ALM: 175 ms, n=9 sessions). Notably, choice probability latencies are not merely reflections of neural activity latencies of these regions (Figure 7B, red bars); while stimulus response latency was earliest in S1, choice probability emerged earliest in wMC.

**Figure 7.**
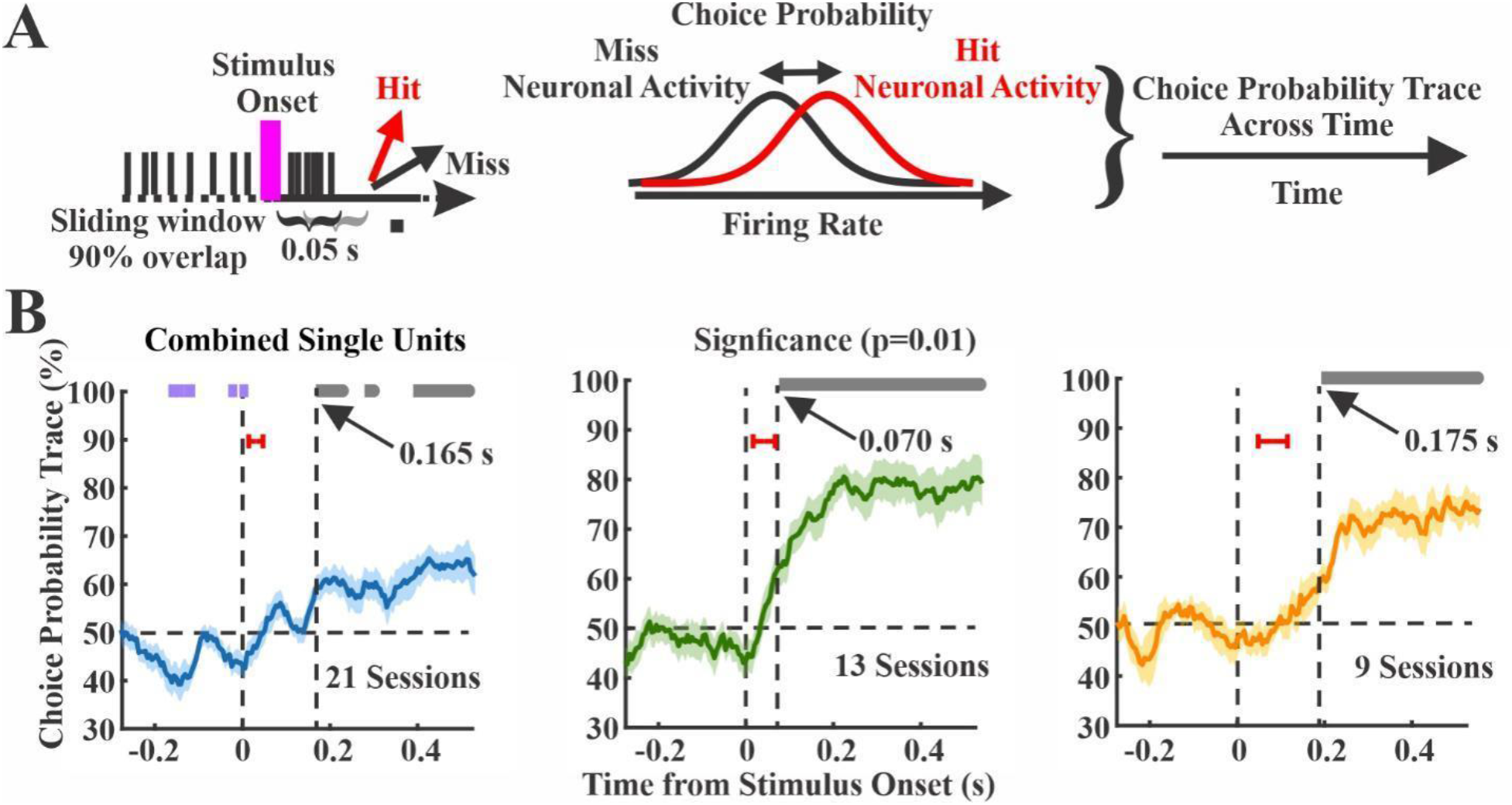
Choice probability within each cortical region. (**A**) A schematic that shows the calculation of the choice probability. Choice probability was calculated by 50 ms sliding window, comparing spike counts on hit (red) versus miss (black) trials. (**B**) Choice probability as a function of time for each region, with overlapping hit and miss distributions at 50% (horizontal dashed line). Data are averages of recording sessions (left, S1, n=21 sessions; middle, wMC, n=13 sessions; right, ALM, n=9 sessions). Significant choice probability is indicated by bars above each plot, gray bars indicate positive choice probability (>50%) whereas purple bars indicate negative choice probability (<50%). Vertical dashed lines indicate latency to significant post-stimulus choice probability. Red bars indicate +/− 1 standard deviation of the sensory response latency for the same sessions. Left, S1 shows pre-stimulus negative choice probability and post-stimulus positive choice probability at a latency of 165 ms. Middle, wMC shows post-stimulus positive choice probability at a latency of 70 ms. Right, ALM shows post-stimulus positive choice probability at 175 ms.

To compare amplitude and time course, in Figure 8 we overlay choice probability signals from all three regions. After stimulus onset, choice probability rises faster in wMC compared to S1 and ALM. We further assessed differences in choice probability magnitude in each time window by conducting pairwise comparisons between regions (Figure 8B). Choice probability was significantly larger during the post-stimulus lockout window in wMC compared to S1 and ALM (two-sample t test, alpha level of 0.05, Bonferroni corrected for the number of regions). Based on these analyses, wMC meets our second criterion for identifying the location of a sensory-motor transformation, in displaying early onset and robust choice probability. Together, our findings support the model presented in Figure 8C.

**Figure 8.**
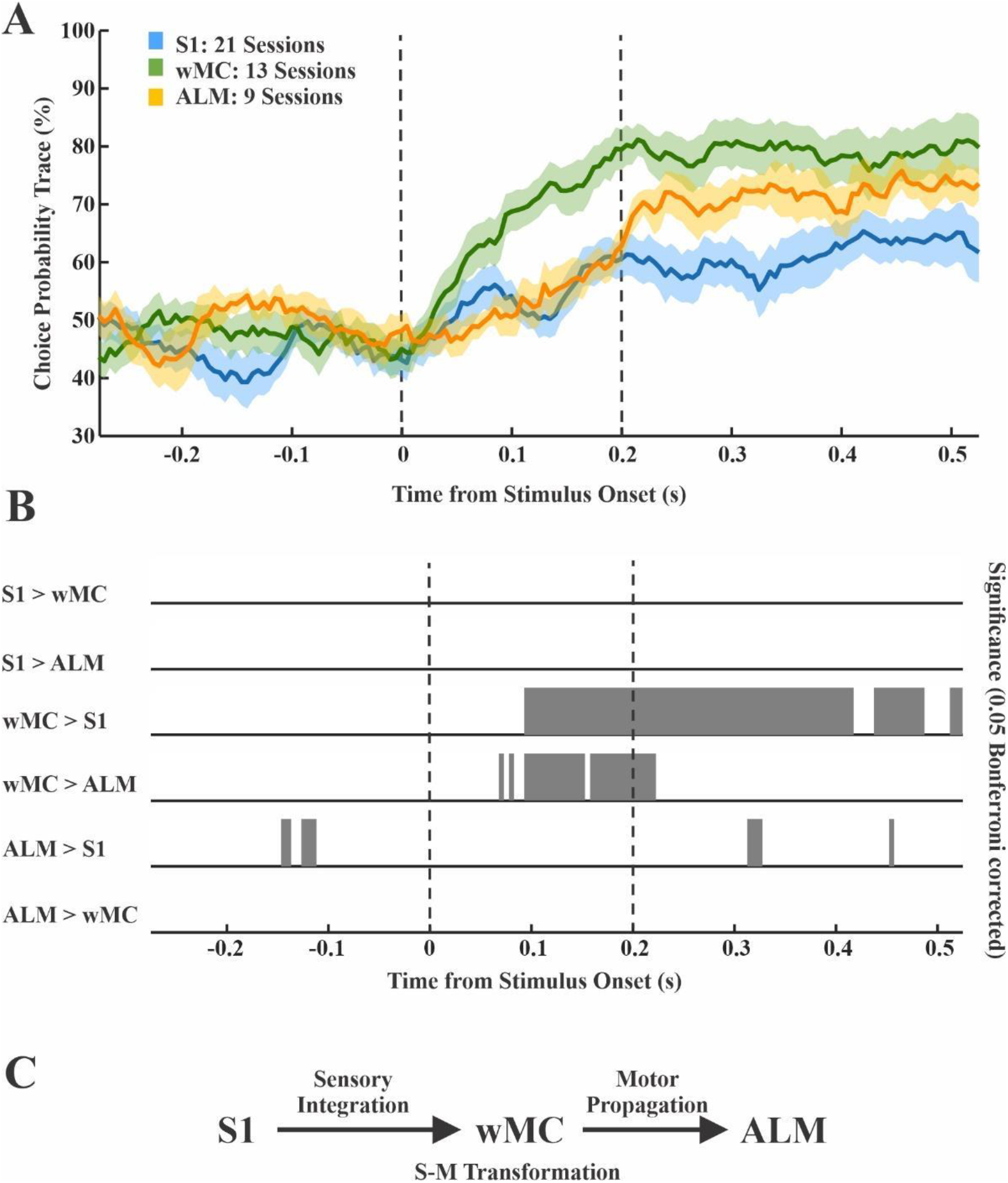
Comparison of choice probability between cortical regions. (**A**) Overlap of traces from Figure 7B. Vertical bars indicate the lockout period, between stimulus onset and start of the response window. Note that wMC rises faster than S1 and ALM and remains elevated throughout the lockout period. (**B**) Results of null hypothesis testing for comparison of choice probability between pairs of regions at each time point. The gray bars denote statistical significance. Choice probability in wMC is greater than S1 and ALM during the lockout period. (**C**) Proposed mechanism: sensory information is integrated from S1 to wMC, the sensory-motor transformation occurs within wMC and a motor signal is propagated to ALM.

## Discussion

The focus of this study is to localize within cortex the region most directly related to the sensory-motor transformation process. This was studied in a whisker detection task, in which mice were trained to respond to a passive whisker deflection by licking a central lickport. Our recordings within cortex focused on three regions which have been identified in a concurrent calcium imaging study (Aruljothi et al., 2020) as potentially contributing to the transformation. Our analyses indicate wMC as the cortical region most directly related to the transformation processes based on having the strongest sensory encoding (Fig 4), robust sensory and motor alignment (Fig 5) and early and robust choice probability (Fig 7 and 8). Our findings are consistent with sensory integration from S1 to wMC, sensory-motor transformation occurring within wMC, followed by propagation of the motor signals to ALM.

The initiation of choice encoding downstream of primary sensory cortices has been demonstrated in studies of non-human primates (Romo et al., 2002; de Lafuente and Romo, 2006; Siegel et al., 2015) and studies of visual detection/discrimination in mouse (Goard et al., 2016; Pho et al., 2018; Salkoff et al., 2020). Our study is also consistent with this finding. However, our study and other studies of the mouse whisker system also show choice encoding in S1 (Sachidhanandam et al., 2013; Yang et al., 2015; Kwon et al., 2016; Aruljothi et al., 2020). Choice encoding in S1 consistently occurs ‘late,’ after the initial feedforward sensory peak activity (Sachidhanandam et al., 2013) (Figure 8). Our findings do not support S1 as initiating the sensory-motor transformation (Figures 7 and 8). We consider two possible causes for S1 choice encoding. First, S1 choice encoding may reflect feedback from choice signals originating in higher order cortices, such as wMC or S2 (Yang et al., 2015; Kwon et al., 2016), as has been described in non-human primates (Siegel et al., 2015). Alternatively, S1 choice probability measurements may not relate to choice encoding at all, but instead reflect re-afferent signals related to the behavioral response sequence. In a related study of the same task, we found whisking to increase approximately 100 ms after stimulus onset, which preceded the onset of licking by approximately 100 ms (Aruljothi et al., 2020). We report here significant choice probability in S1 at 160 ms, 60 ms *after* the onset of whisking. Since whisking is largely absent on miss trials (Aruljothi et al., 2020), re-afferent signals likely contribute to measures of S1 choice probability. In contrast, we find significant choice probability in wMC at 70 ms, 30 ms *before* the onset of whisking. Additionally, we find significant choice probability in ALM at 175 ms, 25 ms before the onset of licking. These neural and behavioral temporal latencies are consistent with the choice-related signals in wMC and ALM initiating the whisking and licking response sequence, respectively.

wMC is a frontal region traditionally studied in the context of whisking initiation and modulation (Carvell et al., 1996; Kleinfeld et al., 1999; Hill et al., 2011). However, recent studies are finding wMC to be an important site of cognitive processing, including sensory integration (Fassihi et al., 2017), motor suppression (Zagha et al., 2015; Ebbesen et al., 2017) and sensory selection (Aruljothi et al., 2020). This study proposes an additional function of sensory-motor transformation, potentially mediated by winner-take-all dynamics in converting a transient, sensory stimulus into a sustained, motor response (Zagha et al., 2015).

We recognize that it is highly unlikely that the sensory-motor transformation occurs exclusively within neocortex. In particular, we suspect that, in our task, interactions between neocortex and striatum are essential for action selection and initiation (Frank, 2011). The question then is, what are the specific contributions of wMC to the transformation process? First, we propose that wMC contributes to sensory integration. An unexpected finding in this study is that sensory encoding is actually enhanced in wMC compared to S1. This finding is based on a larger average neurometric d-prime of wMC neurons and fewer wMC neurons required for the neurometric d-prime to match the psychometric d-prime of the same behavioral sessions (Figure 4). This enhancement may occur by summing the spiking activity of random sets of S1 neurons, as simulated in our pooling analysis. This finding is consistent with a study by Fassihi et al. (2017), in which an extended vibratory stimulus was integrated between S1 and wMC (Fassihi et al., 2017). Interestingly, in that study, perceptual responses matched activity in wMC rather than S1. Thus, a general function of wMC may be to integrate whisker sensory responses. The integrated sensory representations within wMC, rather than S1, may reflect the ‘decision variables’ that ultimately drive behavior (Gold and Shadlen, 2007).

Second, we propose that wMC contributes to response suppression. If wMC were the only transformation node, then wMC suppression ought to prevent whisker stimuli from evoking licking responses (resulting in decreased hit rates). In contrast, in whisker detection tasks, wMC suppression causes variable effects on hit rate, but significantly increases false alarm rates (Huber et al., 2012; Zagha et al., 2015). In the Zagha et al., 2015 study, the false alarms were responses to auditory distractor stimuli, which did not activate wMC. This suggests that wMC may contribute a tonic, global response suppression. One possible organization is that wMC interacts with the basal ganglia (or other subcortical structures) to condition action initiation on whisker detection. Interestingly, in tasks involving different modalities, studies have found suppression of nearby frontal cortex regions to also increase false alarm rates (Narayanan and Laubach, 2006; Goard et al., 2016; Kamigaki and Dan, 2017), suggesting a general function of response suppression for motor cortices (Ebbesen and Brecht, 2017). Much more research is needed to understand how local interactions within wMC and long-range interactions between wMC and subcortical structures mediate sensory selection and the transformation process.

If the cortical transformation does indeed occur within wMC, this has some important implications for sensory-motor transformation in general. First, it would strongly predict that the site of transformation is sensory modality specific. This region of frontal cortex is uniquely connected to whisker sensory cortex compared to other sensory cortices (Mao et al., 2011), and therefore would be unlikely to contribute to the sensory-motor transformation for different sensory modalities. Second, it would predict that the site of transformation may be independent of the ultimate motor response. While there are active debates about the role of wMC in whisker motor control [e.g. (Gao et al., 2003; Haiss and Schwarz, 2005; Matyas et al., 2010; Ebbesen and Brecht, 2017)], this region is not believed to be involved in licking behavior. By comparing neural activity in the same regions during behavioral tasks with different sensory and motor requirements, each prediction may be tested directly.

## Acknowledgements

We thank Dr. Hongdian Yang, Krista Marrero and Krithiga Aruljothi for many helpful discussions throughout the project. This project was supported by the Whitehall Foundation (Research Grant 2017-05-71 to E.Z.) and the National Institutes of Health (R01NS107599 to E.Z.).

